## Supplementary Material for "The interaction between endogenous GABA, functional connectivity and behavioral flexibility is critically altered with advanced age"

Kirstin-Friederike Heise

Movement Control and Neuroplasticity Research Group

KU Leuven

Tervuurse Vest 101; 3001 Leuven, Belgium

### Index

[Supplementary Table 6	Results for logistic GLMM predicting failed transitions [trials with 100% error rate]	6](#_Toc66786173)

[Supplementary Table 7	Results for logistic GLMM to predict fully correct transitions [trials with 0% error]	8](#_Toc66786174)

[Supplementary Table 8	Results for beta GLMM to predict cumulative error rate [0<error rate/100 <1]	9](#_Toc66786175)

[Supplementary Table 12	Rayleigh test for distribution of phase angle differences between left M1 and right M1 sources at baseline [START CUE – 300ms] accounting for GABA+ concentration relative to within group median	14](#_Toc66786186)

[Supplementary Table 13	2-way ANOVA testing GROUP (older vs. young) x GABA(low vs. high) for mean phase angle difference between left M1 and right M1 sources at baseline [START CUE – 300ms]	14](#_Toc66786187)

[Supplementary Figure 3	Association between band-specific M1-M1 phase difference at BASELINE [START CUE – 300ms] and subsequent performance pooled over transition conditions (group average of single trial baseline, error represents subsequent trial following the START CUE)	15](#_Toc66786189)

### Supplementary results for GABA+ data

#### Supplementary Table 1 Descriptive statistics of GABA+ and quantitative quality metrics

|  |  | *LM1* | | | *RM1* | | | *OCC* | | |
| --- | --- | --- | --- | --- | --- | --- | --- | --- | --- | --- |
|  | *Metric* | *mean* | *sd* | *range* | *mean* | *sd* | *range* | *mean* | *sd* | *range* |
| *YOUNG* | GABA+ | 2.517 | 0.244 | 0.91 (2.02-2.93) | 2.389 | 0.241 | 1.17 (1.97-3.13) | 2.859 | 0.23 | 0.82 (2.38-3.2) |
|  | GABA SNR | 31.438 | 5.323 | 21.26 (20.86-42.13) | 28.976 | 3.914 | 13.82 (24.07-37.89) | 23.945 | 5.142 | 23.4 (14.18-37.58) |
|  | GABA Fit Error | 3.564 | 0.802 | 3.04 (2.5-5.54) | 3.94 | 0.69 | 2.77 (2.78-5.55) | 3.861 | 0.58 | 2.48 (2.32-4.8) |
|  | GABA FWHM | 19.728 | 1.088 | 4.68 (17.14-21.82) | 19.852 | 1.428 | 6.06 (15.95-22.01) | 21.104 | 1.03 | 3.56 (19.39-22.95) |
|  | NAA SNR | 350.501 | 69.802 | 240.24 (245.16-485.4) | 323.816 | 118.315 | 391.46 (109.77-501.23) | 360.697 | 75.164 | 250.33 (273.97-524.3) |
|  | Drift | 0.39 | 0.118 | 0.5 (0.22-0.72) | 0.37 | 0.087 | 0.3 (0.21-0.51) | 0.559 | 0.187 | 0.59 (0.27-0.86) |
|  | Frequency Offset | 0.006 | 0.008 | 0.03 (-0.01-0.02) | 0.017 | 0.013 | 0.05 (-0.01-0.04) | 0.002 | 0.004 | 0.02 (0-0.01) |
|  | NAA FWHM | 9.342 | 1.386 | 4.58 (7.53-12.1) | 8.429 | 1.222 | 5.13 (6.38-11.5) | 9.917 | 1.302 | 5.84 (8.24-14.07) |
|  | GM fraction | 0.355 | 0.028 | 0.11 (0.3-0.41) | 0.379 | 0.026 | 0.09 (0.33-0.43) | 0.642 | 0.031 | 0.11 (0.59-0.7) |
|  | WM fraction | 0.562 | 0.03 | 0.12 (0.52-0.63) | 0.523 | 0.028 | 0.1 (0.47-0.57) | 0.271 | 0.032 | 0.13 (0.21-0.34) |
|  | CSF fraction | 0.083 | 0.019 | 0.07 (0.04-0.11) | 0.098 | 0.019 | 0.07 (0.06-0.13) | 0.087 | 0.018 | 0.06 (0.06-0.12) |
| *OLDER* | GABA+ | 2.325 | 0.247 | 1.11 (1.98-3.09) | 2.197 | 0.323 | 1.08 (1.65-2.73) | 2.851 | 0.391 | 1.92 (1.99-3.91) |
|  | GABA SNR | 25.612 | 3.556 | 11.17 (20.1-31.26) | 23.437 | 3.638 | 15.17 (17.4-32.57) | 20.139 | 4.13 | 18.39 (14.06-32.45) |
|  | GABA Fit Error | 3.985 | 0.69 | 2.67 (2.7-5.37) | 4.268 | 1.983 | 8.44 (2.32-10.76) | 4.488 | 1.426 | 5.83 (2.35-8.18) |
|  | GABA FWHM | 20.069 | 1.083 | 4.59 (18.71-23.3) | 19.617 | 1.122 | 4.35 (17.98-22.33) | 21.828 | 1.353 | 5.91 (17.74-23.65) |
|  | NAA SNR | 285.801 | 63.379 | 244.38 (155.31-399.69) | 310.188 | 105.774 | 472.95 (126.56-599.51) | 275.062 | 44.029 | 165.07 (192.78-357.84) |
|  | Drift | 0.632 | 0.308 | 1.39 (0.34-1.73) | 0.677 | 0.236 | 0.9 (0.35-1.26) | 0.816 | 0.268 | 0.79 (0.4-1.19) |
|  | Frequency Offset | 0.007 | 0.01 | 0.05 (-0.01-0.04) | 0.018 | 0.015 | 0.07 (-0.02-0.05) | 0.011 | 0.009 | 0.04 (0-0.04) |
|  | NAA FWHM | 9.647 | 1.012 | 4.05 (8.26-12.31) | 8.789 | 1.04 | 4.03 (6.28-10.31) | 10.114 | 1.226 | 4.7 (7.46-12.16) |
|  | GM fraction | 0.267 | 0.033 | 0.14 (0.2-0.34) | 0.291 | 0.045 | 0.17 (0.2-0.38) | 0.526 | 0.056 | 0.26 (0.36-0.62) |
|  | WM fraction | 0.591 | 0.056 | 0.24 (0.45-0.69) | 0.555 | 0.056 | 0.23 (0.43-0.66) | 0.307 | 0.027 | 0.1 (0.25-0.35) |
|  | CSF fraction | 0.143 | 0.05 | 0.22 (0.06-0.28) | 0.154 | 0.049 | 0.18 (0.09-0.27) | 0.168 | 0.059 | 0.26 (0.08-0.34) |

SNR signal-to-noise ratio, FWHM full width half maximum, GM grey matter, WM white matter, CSF cerebrospinal fluid

#### Supplementary Table 2 Analysis of deviance table for backwards selection of parameters predicting GABA+

| *Parameter* | *X^2^* | *Df* | *p* |
| --- | --- | --- | --- |
| GROUP | 13.467 | 1 | .0002* |
| VOXEL | 44.917 | 2 | <.0001* |
| GABA SNR (centered) | 5.092 | 1 | .024* |
| GABA Fit Error (centered) | 4.007 | 1 | .045* |
| NAA SNR (centered) | 0.199 | 1 | .66 |
| Drift (centered) | 0.128 | 1 | .72 |
| Frequency Offset (centered) | 17.857 | 1 | <.0001* |
| NAA FWHM (centered) | 0.238 | 1 | .63 |
| raw GM fraction (centered) | 15.372 | 1 | <.000* |
| GROUP × VOXEL | 6.942 | 2 | .031* |
| GROUP × GABA SNR (centered) | 0.669 | 1 | .413 |
| VOXEL × GABA SNR (centered) | 0.523 | 2 | .77 |
| GROUP × GABA Fit Error (centered) | 0.408 | 1 | .52 |
| VOXEL × GABA Fit Error (centered) | 9.315 | 2 | .009* |
| GROUP × NAA SNR (centered) | 0.079 | 1 | .78 |
| VOXEL × NAA SNR (centered) | 0.51 | 2 | .78 |
| GROUP × Drift (centered) | 0.491 | 1 | .48 |
| VOXEL × Drift (centered) | 1.03 | 2 | .60 |
| GROUP × Frequency Offset (centered) | 0.007 | 1 | .93 |
| VOXEL × Frequency Offset (centered) | 1.141 | 2 | .57 |
| GROUP × NAA FWHM (centered) | 0.154 | 1 | .70 |
| VOXEL × NAA FWHM (centered) | 3.097 | 2 | .21 |
| GROUP × raw GM fraction (centered) | 8.015 | 1 | .005* |
| VOXEL × raw GM fraction (centered) | 3.411 | 2 | .18 |

#### Supplementary Table 3 Results for Gamma GLMM predicting GABA+ (Final Model)

**Type II Wald statistics**

| *Predictors* | *X^2^* | *df* | *p* |
| --- | --- | --- | --- |
| GROUP | 15.175 | 1 | <.0001 |
| VOXEL | 45.042 | 2 | <.0001 |
| GABA SNR (centered) | 6.736 | 1 | .009 |
| GABA Fit Error (centered) | 5.559 | 1 | .018 |
| raw GM fraction (centered) | 15.363 | 1 | <.0001 |
| NAA SNR (centered) | 0.13 | 1 | .72 |
| Drift (centered) | 0.041 | 1 | .84 |
| Frequency Offset (centered) | 17.197 | 1 | <.0001 |
| NAA FWHM (centered) | 0.123 | 1 | .73 |
| GROUP × VOXEL | 9.566 | 2 | .008 |
| VOXEL × GABA Fit Error (centered) | 5.841 | 2 | .054 |
| GROUP × raw GM fraction (centered) | 6.821 | 1 | .009 |

**Parameter estimates based on references categories indicated**

| *Predictors* | *Estimates (β)* | *std. Error* | *CI* | *Statistic (X^2^)* | *p* |
| --- | --- | --- | --- | --- | --- |
| (Intercept) | 2.855 | 0.264 | 2.337 – 3.373 | 10.799 | **<.0001** |
| Drift (centered) | -0.005 | 0.027 | -0.058 – 0.047 | -0.203 | 0.84 |
| Frequency Offset (centered) | -0.092 | 0.022 | -0.135 – -0.048 | -4.147 | **<.0001** |
| GABA Fit Error (centered) | -0.122 | 0.038 | -0.197 – -0.047 | -3.202 | **.001** |
| GABA SNR (centered) | 0.077 | 0.030 | 0.019 – 0.135 | 2.595 | **.009** |
| raw GM fraction (centered) | -0.031 | 0.154 | -0.334 – 0.272 | -0.201 | .84 |
| NAA FWHM (centered) | 0.009 | 0.025 | -0.039 – 0.057 | 0.351 | .73 |
| NAA SNR (centered) | -0.008 | 0.022 | -0.052 – 0.036 | -0.360 | .72 |
| GROUP [YOUNG] | *Reference* |  |  |  |  |
| GROUP [OLDER] * raw GM fraction | -0.489 | 0.187 | -0.856 – -0.122 | -2.612 | **.009** |
| GROUP [OLDER] * VOXEL [LM1] | -1.190 | 0.388 | -1.950 – -0.430 | -3.070 | **.002** |
| GROUP [OLDER] * VOXEL [RM1] | -1.096 | 0.356 | -1.794 – -0.398 | -3.076 | **.002** |
| GROUP [OLDER] | 0.549 | 0.282 | -0.004 – 1.101 | 1.945 | .052 |
| VOXEL [OCC] | *Reference* |  |  |  |  |
| VOXEL [LM1] * GABA Fit Error (centered) | 0.134 | 0.063 | 0.011 – 0.257 | 2.134 | **.033** |
| VOXEL [LM1] | -0.464 | 0.335 | -1.121 – 0.193 | -1.383 | .17 |
| VOXEL [RM1] | -0.468 | 0.309 | -1.075 – 0.138 | -1.514 | .13 |
| VOXEL [RM1] * GABA Fit Error (centered) | 0.093 | 0.046 | 0.003 – 0.183 | 2.032 | **.04** |
| **Random Effects** | | | | | |
| σ^2^ | 0.01 | | | | |
| τ_00_ _subject_ | 0.01 | | | | |
| ICC | 0.68 | | | | |
| N _subject_ | 44 | | | | |
| Observations | 130 | | | | |
| Marginal R^2^ / Conditional R^2^ | 0.827 / 0.945 | | | | |
| AIC / BIC | -7.34 / 44.27 | | | | |

#### Supplementary Table 4 Model estimated marginal means contrasts for effect of GROUP X VOXEL interaction on GABA+ levels

within group – voxel comparison

| GROUP | VOXEL CONTRAST | Difference | SE | CI_low | CI_high | z | p_holm_ |
| --- | --- | --- | --- | --- | --- | --- | --- |
| OLDER | LM1 -RM1 | -0.09 | 0.065 | -0.28 | 0.101 | -1.382 | 0.52 |
|  | OCC - LM1 | 1.654 | 0.227 | 0.987 | 2.32 | 7.285 | **<.0001** |
|  | OCC - RM1 | 1.564 | 0.215 | 0.933 | 2.196 | 7.271 | **<.0001** |
| YOUNG | LM1-RM1 | 0.005 | 0.071 | -0.204 | 0.213 | 0.065 | .95 |
|  | OCC - LM1 | 0.464 | 0.335 | -0.52 | 1.448 | 1.383 | .52 |
|  | OCC - RM1 | 0.468 | 0.309 | -0.44 | 1.377 | 1.514 | .52 |

between group comparison

| GROUP CONTRAST | VOXEL | Difference | SE | CI_low | CI_high | z | p_holm_ |
| --- | --- | --- | --- | --- | --- | --- | --- |
| YOUNG - OLDER | LM1 | 0.641 | 0.146 | 0.213 | 1.07 | 4.396 | **<.0001** |
|  | RM1 | 0.547 | 0.122 | 0.189 | 0.906 | 4.48 | **<.0001** |
|  | OCC | -0.549 | 0.282 | -1.377 | 0.279 | -1.945 | .26 |

### Supplementary results for behavioural data

#### Supplementary Table 5 Group statistics of outcome parameters of the bimanual transition task

|  | *Total transition* | *Failed*  *transitions* | *Fully correct transitions* | *Cumulative error rate [in %]* | | *Transition latency [in ms]* | |
| --- | --- | --- | --- | --- | --- | --- | --- |
| *GROUP* | *N* | *N (% of total)* | *N (% of total)* | *median* | *ci* | *median* | *ci* |
| young | 2562 | 86 (3.3) | 102 (3.9) | 14.19 | 0.57 | 610 | 10.68 |
| older | 2684 | 198 (7.8) | 285 (10.6) | 14.29 | 0.76 | 807 | 16.77 |

#### Supplementary Table 6 Results for logistic GLMM predicting failed transitions [trials with 100% error rate]

**Type II Wald statistics**

| *Predictors* | *X^2^* | *df* | *p* |
| --- | --- | --- | --- |
| GROUP | 2.19 | 1 | .14 |
| TRANSITION MODE | 34.99 | 1 | <.0001 |
| nTRIALSc | 9.86 | 1 | .002 |
| GROUP × TRANSITION MODE | 1.78 | 1 | .18 |
| GROUP × nTRIALSc | 2.63 | 1 | .11 |
| TRANSITION MODE × nTRIALSc | 0.03 | 1 | .85 |
| GROUP × TRANSITION MODE × nTRIALSc | 4.38 | 1 | .04 |

**Parameter estimates based on references categories indicated**

| *Predictors* | *Odds Ratios* | | *std. Error* | | *CI* | *Statistic (X^2^)* | *p* |
| --- | --- | --- | --- | --- | --- | --- | --- |
| (Intercept)^$^ | -5.66 | | 0.597 | | -6.83 – -4.49 | -9.48 | **<.0001** |
| GROUP[young] | Reference | |  | |  |  |  |
| TRANSITION MODEAP × nTRIALSc | 1.586 | | 0.411 | | 0.954 – 2.635 | 1.78 | .08 |
| GROUP[older] | 2.315 | | 1.772 | | 0.517 – 10.378 | 1.10 | .27 |
| nTRIALSc | 0.699 | | 0.141 | | 0.471 – 1.037 | -1.78 | .08 |
| TRANSITION MODE[into IP] | Reference | |  | |  |  |  |
| GROUP[older] × TRANSITION MODE[into AP] | 1.427 | | 0.472 | | 0.746 – 2.729 | 1.08 | .28 |
| GROUPolder × TRANSITION MODE[into AP] × nTRIALSc | 0.506 | | 0.165 | | 0.267 – 0.957 | -2.10 | **.036** |
| GROUPolder × nTRIALSc | 1.163 | | 0.294 | | 0.709 – 1.908 | 0.60 | .55 |
| TRANSITION MODE[into AP] | 2.075 | | 0.542 | | 1.244 – 3.462 | 2.80 | **.005** |
| **Random Effects** | | | |  |  |  |  |
| σ^2^ | 3.29 | | |  |  |  |  |
| τ_00_ _subjID_ | 4.20 | | |  |  |  |  |
| ICC | 0.56 | | |  |  |  |  |
| N _subjID_ | 42 | | |  |  |  |  |
| Observations | 5124 | | |  |  |  |  |
| Marginal R^2^ / Conditional R^2^ | 0.070 / 0.592 | | |  |  |  |  |
| AIC / BIC | 1329.3 / 1388.1 | | |  |  |  |  |
| ___ |  |  |  |  |  |  |  |

^$^ Intercept given on log scale.

#### Supplementary Table 7 Results for logistic GLMM to predict fully correct transitions [trials with 0% error]

**Type II Wald statistics**

| *Predictors* | *X^2^* | *df* | *p* |
| --- | --- | --- | --- |
| GROUP | 15.426 | 1 | <.0001 |
| TRANSITION MODE | 24.376 | 1 | <.0001 |
| nTRIALSc | 0.046 | 1 | .83 |
| GROUP × TRANSITION MODE | 1.291 | 1 | .26 |
| GROUP × nTRIALSc | 0.567 | 1 | .45 |
| TRANSITION MODE × nTRIALSc | 0.013 | 1 | .91 |
| GROUP × TRANSITION MODE × nTRIALSc | 1.338 | 1 | .25 |

**Parameter estimates based on references categories indicated**

| *Predictors* | *Odds Ratios* | *std. Error* | *CI* | *Statistic (X^2^)* | *p* |
| --- | --- | --- | --- | --- | --- |
| (Intercept)^$^ | -3.26 | 0.277 | -3.80 – -2.72 | -11.77 | **<.0001** |
| GROUP[young] | Reference |  |  |  |  |
| TRANSITION MODE[into AP] × nTRIALSc | 1.264 | 0.283 | 0.816 – 1.959 | 1.05 | .29 |
| GROUP[older] | 3.520 | 1.246 | 1.758 – 7.046 | 3.55 | **<.001** |
| nTRIALSc | 0.979 | 0.124 | 0.764 – 1.254 | -0.17 | .87 |
| TRANSITION MODE[into IP] | Reference |  |  |  |  |
| GROUP[older] × TRANSITION MODE[into AP] | 1.374 | 0.360 | 0.822 – 2.297 | 1.21 | .23 |
| GROUP[older] × TRANSITION MODE[into AP] × nTRIALSc | 0.739 | 0.193 | 0.443 – 1.234 | -1.16 | .25 |
| GROUP[older] × nTRIALSc | 1.009 | 0.153 | 0.749 – 1.359 | 0.06 | .95 |
| TRANSITION MODE[into AP] | 0.443 | 0.099 | 0.286 – 0.688 | -3.63 | **<.001** |
| **Random Effects** | | | | | |
| σ^2^ | 3.29 | | | | |
| τ_00_ _subjID_ | 0.95 | | | | |
| ICC | 0.22 | | | | |
| N _subjID_ | 42 | | | | |
| Observations | 4614 | | | | |
| Marginal R^2^ / Conditional R^2^ | 0.131 / 0.325 | | | | |
| AIC BIC | 2410.4 / 2468.3 | | | | |

^$^ Intercept given on log scale.

#### Supplementary Table 8 Results for beta GLMM to predict cumulative error rate [0<error rate/100 <1]

**Type II Wald statistics**

| *Predictors* | *X^2^* | *df* | *p* |
| --- | --- | --- | --- |
| GROUP | 0.47 | 1 | .49 |
| TRANSITION MODE | 4.909 | 1 | .027 |
| nTRIALSc | 6.692 | 1 | .01 |
| GROUP × TRANSITION MODE | 0.017 | 1 | .90 |
| GROUP × nTRIALSc | 1.18 | 1 | .28 |
| TRANSITION MODE × nTRIALSc | 1.023 | 1 | .31 |
| GROUP × TRANSITION MODE × nTRIALSc | 1.053 | 1 | .31 |

**Parameter estimates based on references categories indicated**

| *Predictors* | *Estimates (β)* | *std. Error* | *CI* | *Statistic (X^2^)* | *p* |
| --- | --- | --- | --- | --- | --- |
| (Intercept) | 0.209 | 0.036 | 0.149 – 0.293 | -9.07 | **<.0001** |
| GROUP[young] | Reference |  |  |  |  |
| TRANSITION MODE[into AP] × nTRIALSc | 0.955 | 0.031 | 0.897 – 1.017 | -1.44 | .15 |
| GROUP[older] | 1.182 | 0.282 | 0.740 – 1.888 | 0.70 | .48 |
| nTRIALSc | 1.067 | 0.024 | 1.021 – 1.116 | 2.87 | **.004** |
| TRANSITION MODE[into IP] | Reference |  |  |  |  |
| GROUP[older] × TRANSITION MODE[into AP] | 0.992 | 0.047 | 0.904 – 1.088 | -0.17 | .87 |
| GROUP[older] × TRANSITION MODE[into AP] × nTRIALSc | 1.049 | 0.049 | 0.957 – 1.150 | 1.03 | .31 |
| GROUP[older] × nTRIALSc | 0.952 | 0.031 | 0.892 – 1.015 | -1.49 | .14 |
| TRANSITION MODE[into AP] | 1.059 | 0.034 | 0.994 – 1.127 | 1.78 | .08 |
| **Random Effects** | | | | | |
| σ^2^ | 0.27 | | | | |
| τ_00_ _subjID_ | 0.59 | | | | |
| ICC | 0.68 | | | | |
| N _subjID_ | 42 | | | | |
| Observations | 4227 | | | | |
| Marginal R^2^ / Conditional R^2^ | 0.010 / 0.687 | | | | |
| AIC / BIC | -7602.95 / -7539.5 | | | | |

^$^ Parameter estimates’ effect on cumulative error rate are given as change in ratio of proportion [exp(logit)].

#### Supplementary Table 9 Results for gamma GLMM to predict transition latency

**Type II Wald statistics**

| *Predictors* | *X^2^* | *df* | *p* |
| --- | --- | --- | --- |
| GROUP | 37.739 | 1 | <.0001 |
| TRANSITION MODE | 8.917 | 1 | <.003 |
| nTRIALSc | 3.949 | 1 | .047 |
| GROUP × TRANSITION MODE | 0.331 | 1 | .56 |
| GROUP × nTRIALSc | 0.13 | 1 | .72 |
| TRANSITION MODE × nTRIALSc | 0.194 | 1 | .66 |
| GROUP × TRANSITION MODE × nTRIALSc | 0.004 | 1 | .95 |

**Parameter estimates based on references categories indicated**

| *Predictors* | *Estimates (β)* | *std. Error* | *CI* | *Statistic (X^2^)* | *p* |
| --- | --- | --- | --- | --- | --- |
| (Intercept) | 568.73 | 25.06 | 521.68 – 620.03 | 143.98 | **<.0001** |
| GROUP[young] | Reference |  |  |  |  |
| TRANSITION MODE[into AP] × nTRIALSc | 1.01 | 0.04 | 0.93 – 1.10 | 0.27 | .79 |
| GROUP[older] | 1.38 | 0.08 | 1.22 – 1.55 | 5.22 | **<.0001** |
| nTRIALSc | 0.96 | 0.03 | 0.91 – 1.02 | -1.37 | .17 |
| TRANSITION MODE[into IP] | Reference |  |  |  |  |
| GROUP[older] × TRANSITION MODE[into AP] | 1.03 | 0.06 | 0.92 – 1.16 | 0.57 | .57 |
| GROUP[older] × TRANSITION MODE[into AP] × nTRIALSc | 1.00 | 0.06 | 0.89 – 1.13 | 0.06 | .95 |
| GROUP[older] × nTRIALSc | 1.01 | 0.04 | 0.93 – 1.09 | 0.21 | .83 |
| TRANSITION MODE[into AP] | 1.07 | 0.04 | 0.99 – 1.17 | 1.73 | .08 |
| **Random Effects** | | | | | |
| σ^2^ | 0.20 | | | | |
| τ_00_ _subjID_ | 0.00 | | | | |
| ICC | 0.02 | | | | |
| N _subjID_ | 42 | | | | |
| Observations | 4614 | | | | |
| Marginal R^2^ / Conditional R^2^ | 0.128 / 0.147 | | | | |
| AIC / BIC | 1339.98 / 1404.31 | | | | |

^$^Parameter estimates’ effect on transition latency and CI given as change in ratio of proportion[exp(logit)].

#### Supplementary Note 1 Thumb reaction time (tRT) – Operationalization and Statistical analysis

Thumb reaction was computed as the response time latency (in ms) between the visual cue occurrence and the respective correct button press for the simple reaction time task. Thumb RTs were trimmed with a lower cut-off at 100ms based on the assumption that a RT <100ms could unlikely be a true reaction to the visual cue and a high cut-off at within group mean+3SD. This conservative approach was chosen to retain as much data as possible without a priori assumptions regarding outlier features.

Thumb RT was optimally fit with a gamma distribution and therefore a GLMM (Gamma family with identity link) was fitted to predict tRT with GROUP, SIDE, and nTRIALSc.. Factors GROUP (old, young), SIDE (left, right), and covariate nTRIALSc (trial number, centred) were entered as fixed effects. Random intercepts were fit on subject level. Trimming of tRT data resulted in discarding 1.8% of trials resulting in a total of 1176 tRT trials entering the analysis.

The model's total explanatory power is substantial (conditional R2 = 0.80) and the part related to the fixed effects alone (marginal R2) is of 1.0. The model's intercept, corresponding to GROUP = young, side= right, nTRIALSc = 0, is at 501.0ms (SE = 18.17, 95% CI [465.39, 536.61], p < .0001). Older participants were on average over 252ms slower as shown by the positive effect of GROUP[older] (beta = 252.70±16.38, 95% CI [220.59, 284.80], p < .0001). Overall, left side responses were more than 28ms slower compared to right side responses (beta = 28.89±9.21, 95% CI [10.84, 46.95], p < .01). A positive interaction effect of SIDE [left] on GROUP [older] (beta = 32.15±14.38, 95% CI [3.97, 60.34], p < .05) was followed up by contrasts estimated from marginal means, which revealed that both groups responded significantly slower with the left hand (OLDER| right – left: ΔEMM = -61.047± 13.909, 95% CI [-97.741, -24.353], z = -4.389, p<.0001; YOUNG| right – left: ΔEMM = -28.893± 9.213, 95% CI [-53.199, -4.587], z = -3.136, p <.01). Time across the experiment had neither a main effect on tRT (beta = -8.77±6.36, 95% CI [-21.23, 3.69], p >.1), nor was a relevant effect modulator of the other factors.

#### Supplementary Table 10 Results for gamma GLMM to predict thumb reaction time

**Parameter estimates based on references categories indicated**

| *Parameter* | *Estimates (β)* | *std. Error* | *CI* | *Statistic (X^2^)* | *p* |
| --- | --- | --- | --- | --- | --- |
| (Intercept) | 501.00 | 18.17 | 465.39 – 536.61 | 27.58 | <.0001 |
| nTRIALSc | -8.77 | 6.36 | -21.23 – 3.69 | -1.38 | .17 |
| GROUP[young] | *Reference* |  |  |  |  |
| GROUP[older] x nTRIALSc | -10.84 | 9.01 | -28.51 – 6.82 | -1.20 | .23 |
| GROUP[older] x SIDE[left] | 32.15 | 14.38 | 3.97 – 60.34 | 2.24 | **.03** |
| GROUP[older] x SIDE[left] x nTRIALSc | -3.36 | 11.67 | -26.22 – 19.51 | -0.29 | .77 |
| GROUP[older] | 252.70 | 16.38 | 220.59 – 284.80 | 15.42 | **<.0001** |
| SIDE[right] | *Reference* |  |  |  |  |
| SIDE[left] x nTRIALSc | 6.75 | 9.74 | -12.34 – 25.85 | 0.69 | .49 |
| SIDE[left] | 28.89 | 9.21 | 10.84 – 46.95 | 3.14 | **.002** |
| *Random Effects* | | | | | |
| σ^2^ | 0.07 | | | | |
| τ_00_ _subject_ | 4496.13 | | | | |
| ICC | 1.00 | | | | |
| N _subject_ | 43 | | | | |
| Observations | 1176 | | | | |
| Marginal R^2^ / Conditional R^2^ | 0.80 / 1.0 | | | | |
| AIC / BIC | 15437.25 /15487.95 | | | | |

### Supplementary material for the EEG data analysis

#### Supplementary Note 2 Statistical analysis of ask related spectral power changes

First, task-related power change, dB, from baseline was analyzed using a cluster corrected permutation (1000 permutations, 2-tailed t-test, p<.05) within subject to extract effect size of change from baseline irrespective of transition mode. This step was used to extract the z-transformed power changes per condition within subject. Second, group level cluster-based permutation analysis (1000 permutations, 2-tailed t-test, p<.05) of change in power from baseline pooled over transition modes (stimulus-locked analysis) was used to confirm the relevance of spectral modulation within the selected time and frequency window. Third, in order to test the effect transition mode and its modulation by age group, differences of the z matrices were calculated for the transition mode contrast (IP – AP) for the age groups separately and subsequently subtracted from each other ([IP-AP]YOUNG – [IP-AP]OLDER). A two-sided t-test (p<.05) was then run with permuting the age group allocation (1000 permutations). The third step was performed relative to the response, i.e. ±260ms around the individual median transition latency specific for IP and AP transitions, respectively (response-locked analysis). In all three steps, clusters were corrected for multiple comparison and considered significant if they contained more time × frequency data points than expected under the null hypothesis at p<.05.

#### Supplementary Figure 1 Statistical results of spectral power changes.

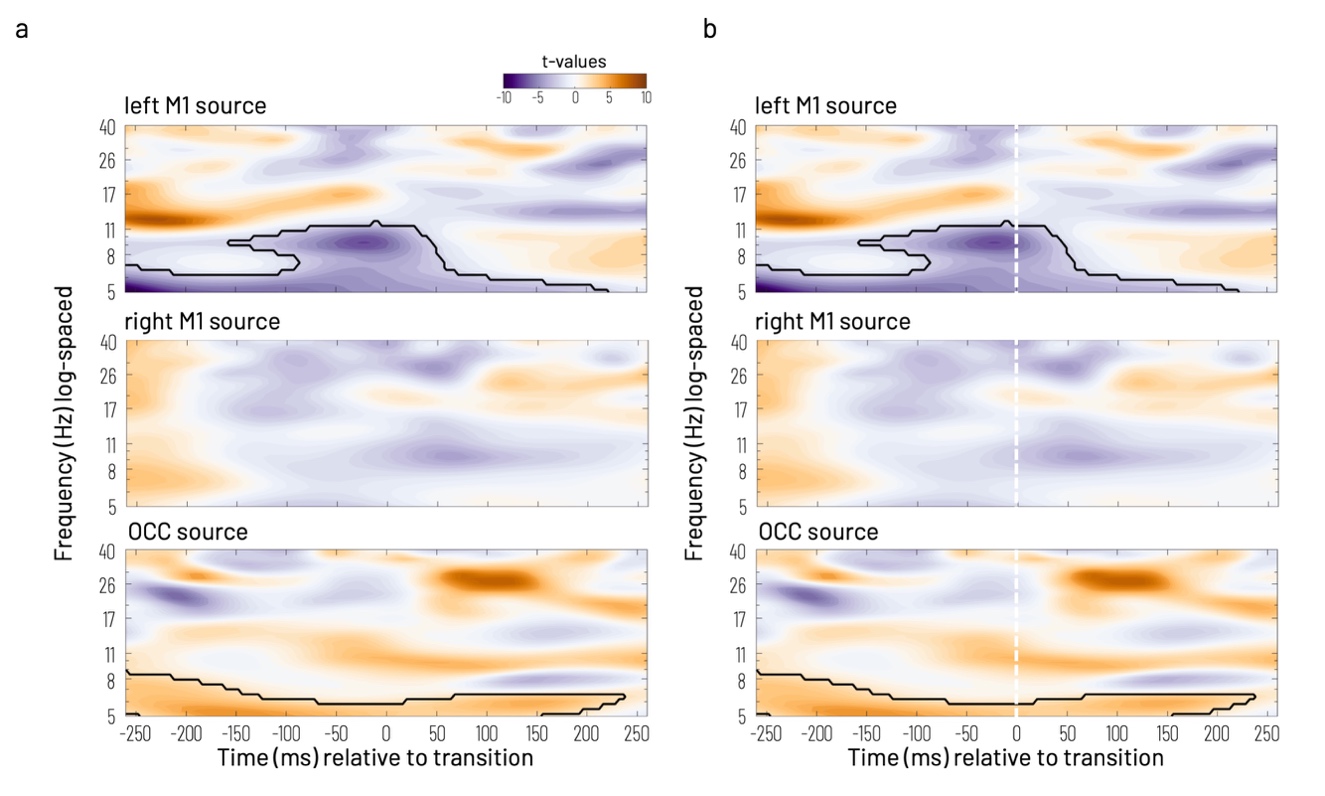

**Supplementary Figure 2 a)**	**Statistical results of cluster corrected permutation test for significant power changes from baseline pooled over age groups and transition modes.** Time (in sec) is presented relative to visual cue onset (indicated by black arrows). Orange colour shading indicates significant power increase and purple colour shading indicates significant power decrease with respect to the baseline period [-500 to -200ms relative to cue onset]. Black lines highlight significant frequency-by-time clusters (2-sided t-test, permutation-based cluster correction). **b)**	**Statistical results of cluster corrected permutation test for significant power changes for group contrast (OLDER vs. YOUNG) for the transition mode difference (IP - AP).** Orange colour shading indicates significant power increase and purple colour shading indicates significant power decrease for the OLDER relative to the YOUNG. Black lines highlight significant frequency-by-time clusters (2-sided t-test, permutation-based cluster correction at p<.05). Time (in ms) is presented relative to the response (white vertical dashed line at 0ms). Zooming into the time window ±260ms around the individual median transition latency for the analysis of the effect of transition mode and its modulation by factor age group, i.e. running a two samples t-test on the contrast [IP-AP]YOUNG – [IP-AP]OLDER, revealed specific time and frequency clusters reaching level of significance (p<.05, 2-sided) for the individual sources.

#### Supplementary Figure 2 Stimulus-locked analysis of ISPC modulation.

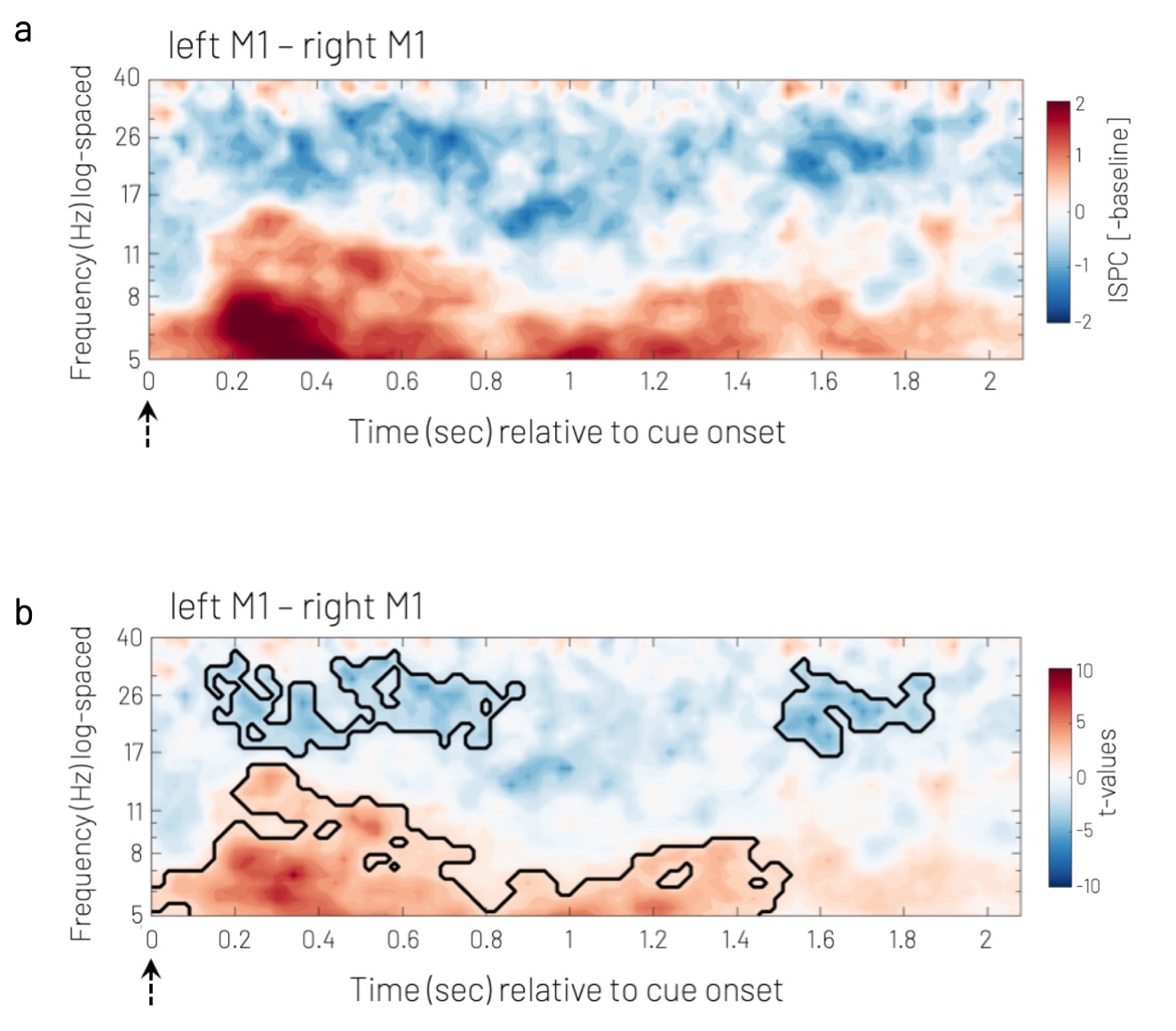

**Supplementary Figure 3a)	Baseline subtracted ISPC.** Time and frequency distribution of change in connectivity to baseline (-500 to -200ms before visual cue onset) for left M1 – right M1 sources pooled over age groups and transition modes. Red shading indicates a relative increase in connectivity, blue shading indicates a relative decrease. Black arrow at time 0ms indicates visual start cue. **b)** **Statistical results of time and frequency resolved modulation of connectivity** relative to baseline (ISPC baseline subtracted). Clusters of significant connectivity change (corrected t-values shown for p<.05, 2-tailed) are highlighted with black lines. Colour coding of direction of effects as in a). The analysis of connectivity modulation revealed significant time × frequency clusters when pooled over age groups and transition modes. The left M1 – right M1 connection showed a distributed connectivity increase in the theta, alpha, and mu range from cue onset until 1500ms. Connectivity increased most prominently around 300ms in the theta and low alpha range. Around this time, there was also increased connectivity in the upper alpha/mu and low beta range, which was not visible later in the time window of interest. M1-M1 connectivity significantly decreased in the beta spectrum (17 – 40 Hz) around 200ms until 900ms following the visual cue onset. A second cluster between 1500 until 1900ms showed also a connectivity decrease in the beta frequency band.

### Supplementary results for phase angle differences

#### Supplementary Note 3 Rayleigh test for distribution of phase angle differences between left M1 and right M1 sources at time of transition pooled over groups and pooled over transition modes.

Distribution of phase angle differences between left and right M1 sources was non-uniform for the young in the low beta range (15-22Hz: z = 70.43, p_FDR_= 1.76e-30) and for both age groups in the high beta range (YOUNG: z = 4.69, p_FDR_ = .03, OLDER: z=7.26, p_FDR_ = .003) when pooled over transition conditions. When pooled over age groups, non-uniformity of phase angle-differences between left and right M1 sources was shown for transitions into IP for the low beta band only (15-22Hz: z = 11.21, p_FDR_ = 4.99e-05), whereas for transitions into AP this was the case for both, the low and the high beta band (15-22Hz: z = 12.21, p_FDR_ = 2.45e-05; 25-30Hz: z = 4.09, p_FDR_ =.05).

#### Supplementary Table 11 Rayleigh test for distribution of phase angle differences between left M1 and right M1 sources at time of transition accounting for GABA+ concentration relative to within group median

| GABA+ relative to group median | group | frequency range (Hz) | z | p_FDR_ |
| --- | --- | --- | --- | --- |
| low | YOUNG | 15 - 22 | 25.16 | 1.08e-10*** |
| low | YOUNG | 25 - 30 | 2.87 | 0.21 |
| low | OLDER | 15 - 22 | 2.66 | 0.24 |
| low | OLDER | 25 - 30 | 6.51 | 0.008* |
| high | YOUNG | 15 - 22 | 47.58 | 2.53e-20*** |
| high | YOUNG | 25 - 30 | 6.38 | 0.008* |
| high | OLDER | 15 - 22 | 1.42 | 0.75 |
| high | OLDER | 25 - 30 | 5.56 | 0.02* |

#### Supplementary Table 12 Rayleigh test for distribution of phase angle differences between left M1 and right M1 sources at baseline [START CUE – 300ms] accounting for GABA+ concentration relative to within group median

| GABA+ relative to group median | group | frequency range (Hz) | z | p_FDR_ |
| --- | --- | --- | --- | --- |
| low | YOUNG | 15 - 22 | 55.51 | 8.85e-24 |
| low | YOUNG | 25 - 30 | 6.60 | 0.006 |
| low | OLDER | 15 - 22 | 47.29 | 2.05e-20 |
| low | OLDER | 25 - 30 | 8.29 | 0.001 |
| high | YOUNG | 15 - 22 | 81.41 | 6.86e-35 |
| high | YOUNG | 25 - 30 | 5.34 | 0.02 |
| high | OLDER | 15 - 22 | 12.95 | 1.47e-05 |
| high | OLDER | 25 - 30 | 2.27 | n.s. (0.32) |

#### Supplementary Table 13 2-way ANOVA testing GROUP (older vs. young) x GABA(low vs. high) for mean phase angle difference between left M1 and right M1 sources at baseline [START CUE – 300ms]

| Frequency range | Source | d.f. | X^2^ | P-Value |
| --- | --- | --- | --- | --- |
| 15-22Hz | GROUP | 2 | 71.09 | 3.33e-16 |
|  | GABA+ | 2 | 47.87 | 4.04e-11 |
|  | Interaction | 1 | 87.38 | 0 |
| 25-30Hz | GROUP | 2 | 5.10 | 0.05 |
|  | GABA+ | 2 | 10.25 | 0.006 |
|  | Interaction | 1 | 8.40 | 0.004 |

#### Supplementary Table 14 Circular-linear correlation phase angle difference ~ subsequent error

| GABA+ relative to group median | group | frequency range (Hz) | rho | p_FDR_ |
| --- | --- | --- | --- | --- |
| low | YOUNG | 15 - 22 | 0.05 | 0.0001 |
| low | YOUNG | 25 - 30 | 0.01 | n.s. (2.40) |
| low | OLDER | 15 - 22 | 0.10 | 0 |
| low | OLDER | 25 - 30 | 0.08 | 3.61e-06 |
| high | YOUNG | 15 - 22 | 0.11 | 0 |
| high | YOUNG | 25 - 30 | 0.10 | 5.28e-08 |
| high | OLDER | 15 - 22 | 0.06 | 6.10e-07 |
| high | OLDER | 25 - 30 | 0.02 | n.s. (1.09) |

#### Supplementary Figure 3 Association between band-specific M1-M1 phase difference at BASELINE [START CUE – 300ms] and subsequent performance pooled over transition conditions (group average of single trial baseline, error represents subsequent trial following the START CUE)

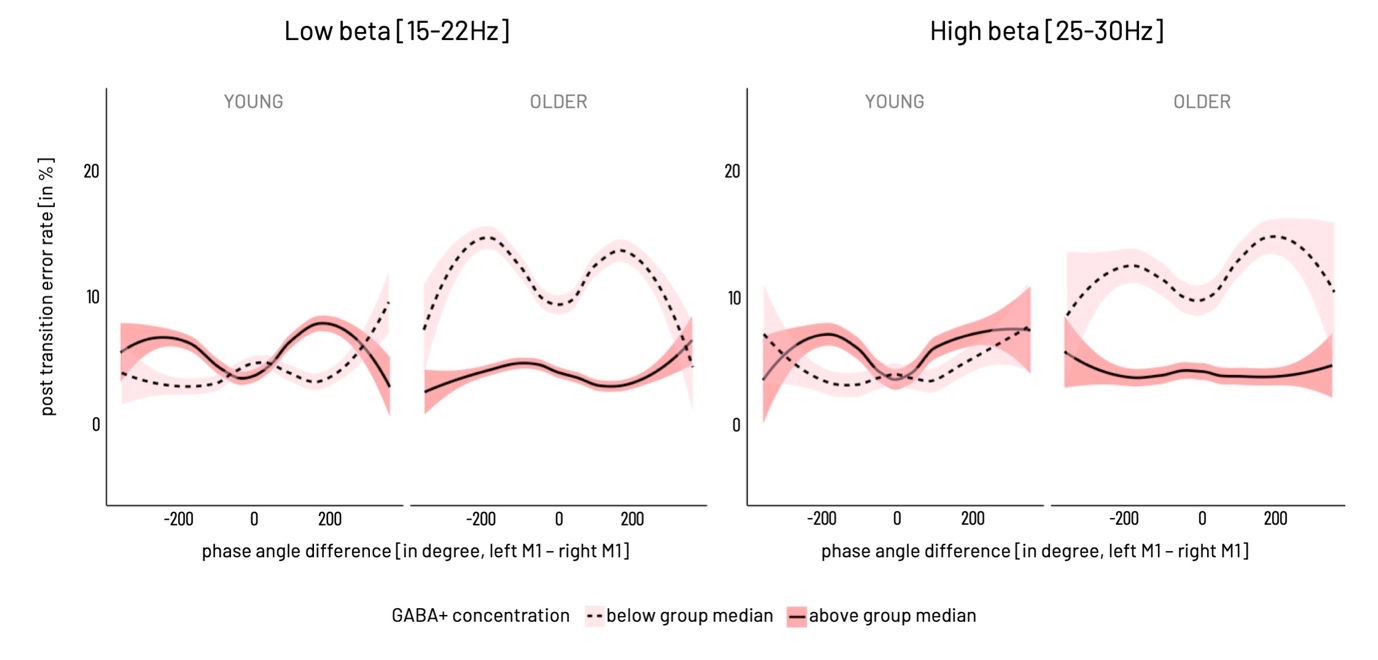

#### Supplementary Table 15 Comparison M1M1 – OCC-M1

Rayleigh test for non-uniformity of phase angle difference distribution

OCC-LM1

| Interaction between | GABA+ relative to group median | group | frequency range (Hz) | z | p_FDR_ |
| --- | --- | --- | --- | --- | --- |
| OCC-LM1 | low | YOUNG | 15 - 22 | 16.88 | 1.73e-06 |
|  | low | YOUNG | 25 - 30 | 0.73 | 1.81 |
|  | low | OLDER | 15 - 22 | 5.27 | 0.05 |
|  | low | OLDER | 25 - 30 | 2.27 | 0.68 |
|  | high | YOUNG | 15 - 22 | 1.30 | 1.30 |
|  | high | YOUNG | 25 - 30 | 6.21 | 0.023 |
|  | high | OLDER | 15 - 22 | 0.72 | 1.81 |
|  | high | OLDER | 25 - 30 | 0.34 | 2.42 |
| OCC -RM1 | low | YOUNG | 15 - 22 | 1.0 | 1.37 |
|  | low | YOUNG | 25 - 30 | 0.11 | 2.79 |
|  | low | OLDER | 15 - 22 | 8.96 | 0.002 |
|  | low | OLDER | 25 - 30 | 6.17 | 0.02 |
|  | high | YOUNG | 15 - 22 | 4.14 | 0.07 |
|  | high | YOUNG | 25 - 30 | 11.21 | 0.0003 |
|  | high | OLDER | 15 - 22 | 0.83 | 1.47 |
|  | high | OLDER | 25 - 30 | 5.19 | 0.03 |

#### Supplementary Table 16 2-way ANOVA testing mean direction CONNECT (LM1 -RM1 vs. OCC-LM1) x group (older vs young)

| Frequency range | Source | d.f. | X^2^ | P-Value |
| --- | --- | --- | --- | --- |
| 15-22Hz | CONNECT | 2 | 10.78 | 0.005 |
|  | GROUP | 2 | 50.07 | 1.34e-11 |
|  | Interaction | 1 | 51.73 | 6.36e-13 |
| 25-30Hz | CONNECT | 2 | 0.70 | 0.70 |
|  | GROUP | 2 | 7.15 | 0.03 |
|  | Interaction | 1 | 15.40 | 8.72e-05 |

#### Supplementary Table 17 2-way ANOVA testing mean direction CONNECT (LM1 -RM1 vs. OCC-RM1) x GROUP (older vs young)

**15-22Hz**

| Frequency range | Source | d.f. | X^2^ | P-Value |
| --- | --- | --- | --- | --- |
| 15-22Hz | CONNECT | 2 | 13.01 | 0.002 |
|  | GROUP | 2 | 43.67 | 3.30e-10 |
|  | Interaction | 1 | 56.10 | 6.90e-14 |
| 25-30Hz | CONNECT | 2 | 6.76 | 0.03 |
|  | GROUP | 2 | 41.65 | 9.04e-10 |
|  | Interaction | 1 | 9.67 | 0.002 |

#### Supplementary Table 18 Circular-linear correlation phase angle difference ~ subsequent error

| Interaction between | GABA+ relative to group median | group | frequency range (Hz) | rho | p_FDR_ |
| --- | --- | --- | --- | --- | --- |
| OCC – LM1 | low | YOUNG | 15 - 22 | 0.024 | ns (0.44) |
|  | low | YOUNG | 25 - 30 | 0.04 | ns (0.36) |
|  | low | OLDER | 15 - 22 | 0.03 | ns (0.14) |
|  | low | OLDER | 25 - 30 | 0.04 | ns (0.21) |
|  | high | YOUNG | 15 - 22 | 0.03 | ns (0.36) |
|  | high | YOUNG | 25 - 30 | 0.04 | ns (0.43) |
|  | high | OLDER | 15 - 22 | 0.01 | ns (1.88) |
|  | high | OLDER | 25 - 30 | 0.05 | ns (0.14) |
| OCC – RM1 | low | YOUNG | 15 - 22 | 0.02 | ns (1.46) |
|  | low | YOUNG | 25 - 30 | 0.05 | ns (0.62) |
|  | low | OLDER | 15 - 22 | 0.02 | ns (0.69) |
|  | low | OLDER | 25 - 30 | 0.01 | ns (2.87) |
|  | high | YOUNG | 15 - 22 | 0.02 | ns (1.19) |
|  | high | YOUNG | 25 - 30 | 0.02 | ns (2.02) |
|  | high | OLDER | 15 - 22 | 0.01 | ns (2.02) |
|  | high | OLDER | 25 - 30 | 0.02 | ns (1.98) |

#### Supplementary Figure 4 Association between band-specific OCC-M1 phase difference at time of transition and subsequent performance pooled over transition conditions

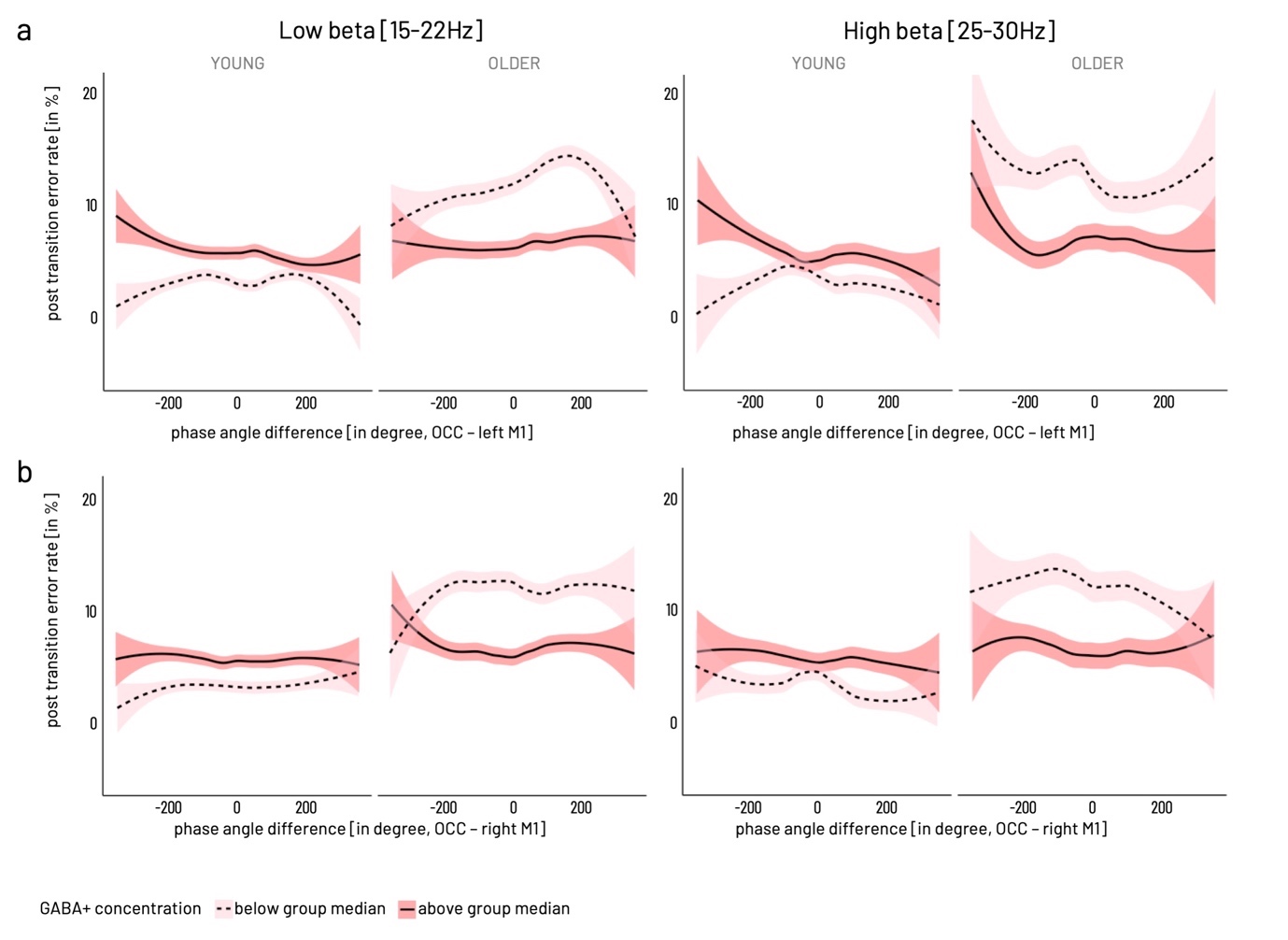

**Supplementary Figure 5 a)** Phase angle difference between occipital and left M1 source (OCC-left M1). **b)** Phase angle difference between occipital and right M1 source (OCC-right M1).

### Supplementary results for the Bayesian moderated mediation analyses

#### Supplementary Table 19 Regression coefficients of Bayesian moderated mediation models

| DV | Mediator | LEFT M1 GABA+ | | | RIGHT M1 GABA+ | | |
| --- | --- | --- | --- | --- | --- | --- | --- |
|  | Time Window | PRE | PERI | POST | PRE | PERI | POST |
|  | Frequency band of IV | high beta | low beta | high alpha / mu | high beta | low beta | high alpha / mu |
| error rate | τ | 0.032  [-0.057, 0.121]  71.5% | **0.087**  **[0.031, 0.141]**  **99.5%**** | **0.104**  **[0.06, 0.145]**  **100%**** | 0.055  [-0.041, 0.151]  81.9% | **0.084**  **[0.026, 0.136]**  **99.2%^*^** | **0.104**  **[0.061, 0.146]**  **99.9%**** |
|  | ⍺ | 0.073  [-0.216, 0.361]  65.7% | **0.447**  **[0.25, 0.65]**  **100%**** | -0.007  [-0.157, 0.128]  53.1% | **-0.507**  **[-0.897, -0.123]**  **98.2%*** | **0.381**  **[0.164, 0.601]**  **99.7%**** | 0.148  [-0.044, 0.345]  88.9% |
|  | β | 0.032  [-0.028, 0.092]  79.8% | 0.028  [-0.019, 0.072]  82.9% | 0.041  [-0.021, 0.099]  85.2% | -0.035  [-0.09, 0.021]  84.5% | **-0.049**  **[-0.086, -0.011]**  **98.2%**** | -0.05  [-0.1, 0.001]  94.4% |
|  | ⍺ × β | 0.001  [-0.011, 0.017]  59.6% | 0.011  [-0.01, 0.033]  82.9% | 0.0  [-0.008, 0.007]  52.4% | 0.015  [-0.012, 0.052]  83.3% | **-0.017**  **[-0.036, -0.001]**  **97.9%*** | -0.006  [-0.02, 0.005]  84.5% |
|  | τ’ | 0.029  [-0.059, 0.117]  70.2% | **0.075**  **[0.017, 0.131]**  **98.1%*** | **0.104**  **[0.062, 0.146]**  **100%**** | 0.037  [-0.056, 0.13]  73.6% | **0.102**  **[0.048, 0.157]**  **99.8%**** | **0.112**  **[0.068, 0.151]**  **100%**** |
| transition latency | τ | -0.005  [-0.172, 0.159]  52.0% | **0.186**  **[0.093, 0.281]**  **99.9%**** | **0.181**  **[0.098, 0.27]**  **100%**** | -0.082  [-0.252, 0.089]  78.2% | **0.138**  **[0.04, 0.235]**  **98.8%*** | **0.143**  **[0.055, 0.232]**  **99.3%*** |
|  | ⍺ | 0.075  [-0.214, 0.356]  66.2% | **0.448**  **[0.253, 0.653]**  **100%**** | -0.007  [-0.154, 0.129]  53.2% | **-0.506**  **[-0.897, -0.125]**  **98.2%*** | **0.381**  **[0.16, 0.595]**  **99.7%**** | 0.148  [-0.041, 0.34]  89.4% |
|  | β | -0.034  [-0.163, 0.085]  67.3% | -0.048  [-0.136, 0.047]  79.8% | -0.003  [-0.123, 0.116]  51.6% | 0.049  [-0.071, 0.168]  74.9% | 0.052  [-0.026, 0.133]  85.3% | 0.055  [-0.055, 0.162]  78.5% |
|  | ⍺ × β | -0.001  [-0.028, 0.021]  55.1% | -0.02  [-0.066, 0.02]  79.8% | 0  [-0.011, 0.009]  50.0% | -0.02  [-0.096, 0.038]  74.1% | 0.018  [-0.013, 0.053]  85.1% | 0.005  [-0.011, 0.033]  72.5% |
|  | τ’ | -0.003  [-0.161, 0.165]  50.9% | **0.208**  **[0.109, 0.303]**  **100%**** | **0.181**  **[0.093, 0.264]**  **100%**** | -0.057  [-0.225, 0.115]  70.8% | **0.118**  **[0.026, 0.217]**  **97.6%*** | **0.135**  **[0.05, 0.221]**  **99.3%*** |

Regression coefficient of model paths given as median [89% HDI], pd (in %). Asterisks indicate approximate 2-tailed p-value ˚ p<.1, * p<.05, **p<.01
